# Individual variability in performance reflects selectivity of the multiple demand network among children and adults

**DOI:** 10.1101/2022.08.11.503644

**Authors:** Elana Schettini, Kelly J. Hiersche, Zeynep M. Saygin

**Affiliations:** Department of Psychology, The Ohio State University, Columbus, OH, USA; Center for Cognitive and Behavioral Brain Imaging, The Ohio State University, Columbus, OH, USA

**Keywords:** multiple demand network, spatial working memory, functional connectivity, cognitive load, child, development, selectivity, fMRI, resting-state

## Abstract

Executive function (EF) is essential for human cognition, allowing individuals to effectively engage in cognitively demanding tasks. In adults, EF is subserved by a set of frontoparietal brain regions (termed the multiple demand (MD) network) which show robust responses to a wide variety of cognitively demanding tasks (i.e., domain-general) and reflect cognitive effort exerted on the task. But while essential, children initially show poor EF skills with prolonged development of these skills. Do children recruit the same network as adults? Is it functionally and connectionally distinct from adjacent language cortex as it is in adults? And is this activation or connectivity dependent on age or on the individual’s EF task performance? We scanned 44 adults and 37 children ages 4-12 years for two separate tasks (MD spatial working memory task and passive language task) and a resting-state fMRI scan. Because motion is a concern in child samples, we asked a subset of adult subjects to participate in additional “wiggly” scans of the MD task and to move slightly during those scans. We defined subject-specific functional regions of interest (ss-fROIs) and found bilateral activation of the MD network in children. In both children and adults, these ss-fROIs are not recruited for linguistic processing and are also connectionally distinct from language ss-fROIs. MD activation in children was lower than that observed in adults, but it was unrelated to motion as evidenced by motion-matched comparisons between children and adult groups and by repeated measures comparisons within the adult group. Right-lateralized ss-fROIs showed increasing load-based MD responses that were robustly associated with performance, demonstrated both cross-sectionally and in a subset of children scanned longitudinally about one year apart. These data suggest that even in young children the MD network is selective to cognitive demand, is distinct from adjacent cortex, and increases in its selectivity as a child improves their EF skills, independently of age. Overall, these findings show that neural structures subserving domain-general EF emerge early and are sensitive to ability in both children and adults. This research advances our understanding of how high-level human cognition emerges and can inform interventions targeting cognitive control.

**Significance statement:** This study provides evidence that young children already show differentiated brain network organization between regions that process cognitive demand and language. These data support the hypothesis that children recruit a similar network as adults to process cognitive demand, and despite immature characteristics, children’s selectivity looks more adult-like as their executive function ability increases. Mapping early stages of network organization furthers our understanding of the functional architecture underlying domain-general executive function. Determining typical variability underlying cognitive processing across developmental periods helps establish a threshold for executive dysfunction. Early markers of risk are necessary for effective early identification, prevention, and intervention efforts for individuals struggling with deficits in processing cognitive demand.

## Introduction

Executive function (EF) is comprised of three distinct, yet intersecting, components (working memory, cognitive shifting, and inhibitory control) that, together, facilitate effective organization, planning, and self-regulation (Friedman & Miyake, 2017; Miyake & Friedman, 2012). Children and adolescents, whose impulse control is still developing, consistently perform worse on laboratory measures of EF compared to adults and overall performance on EF tasks improves with age (Anderson, 2002; Westerberg, Hirvikoski, Forssberg, & Klingberg, 2004; Zelazo, Craik, & Booth, 2004; Best & Miller, 2010; Wiebe & Karbach, 2017). So, do children process cognitively demanding tasks using the same system as adults? Factor analytic studies support a hierarchical structure of EF in adults, with an overarching “common EF” factor (Friedman & Miyake, 2017). However, developmental studies support a unitary model of EF that only starts to dissociate in mid-to-late childhood (Wiebe, Espy, & Charak, 2008; Wiebe et al., 2011; Brydges, Fox, Reid, & Anderson, 2014; Lerner & Lonigan, 2014). Duncan (2010) proposed that the domain-general multiple demand (MD) network, comprised of frontal, parietal, cingular, and opercular brain regions, subserves this “common EF” factor.

Previous work examined the MD network in adults and showed that the MD network is reliably recruited across a variety of EF tasks (Niendam et al., 2012; Fedorenko, Duncan, & Kanwisher, 2013; Shashidhara, Spronkers, & Erez, 2020). Fedorenko and colleagues (2013) identified 10 bilateral regions that show activation in a majority of their adult sample during different types of EF tasks. Further, these core MD regions are distinct from the adjacent language network in both functional specificity (Fedorenko, Duncan, & Kanwisher, 2013; Diachek, Blank, Siegelman, Affourtit, & Fedorenko, 2020) and resting-state connectivity (Blank, Kanwisher, & Fedorenko, 2014). In addition to showing robust activation for any type of cognitive demand, activation of the MD network is correlated with behavioral markers, such as reaction time, accuracy, and intelligence (Assem, M., Blank, Mineroff, Ademoğlu & Fedorenko, 2020). This suggests that these regions are sensitive to EF performance and ability. Thus, we might expect that activation of children’s MD network would reflect their immature EF skills.

It is possible that the neural development of the MD network parallels development of behavioral performance on EF tasks. In this case, we’d expect that as a child’s performance improves, their MD network activation would look more adult-like. Similar patterns of activation are observed across EF tasks among typically developing children and adults, though child studies show more variability in strength and location of MD activation (Vogan, Morgan, Powell, Smith, & Taylor, 2016; McKenna, Rushe, & Woodcock, 2017; Fiske & Holmboe, 2019; Durston et al., 2022). It remains unclear whether any developmental differences in MD network activation reflects age-dependent gains independent of behavioral performance on EF tasks, or whether the activation of this network more closely depends on behavioral performance regardless of age. It is also unclear whether this MD network is functionally and connectionally distinct from adjacent domain-specific language areas in children, like they are in adults.

To date, no one has looked at neural processing of EF in comparison to other mental functions, like language, in a sample of children. In this study, we investigate the cross-sectional and longitudinal development of the MD network 4-12-year-old children. We collected task-dependent functional magnetic resonance imaging (fMRI) to identify brain regions that children recruit while engaging in a cognitively demanding spatial working memory (SWM) task. We also collected a high-level language processing task and a resting-state scan on these children. To account for stable individual variability in anatomical and functional network organization (Gratton et al., 2018), which is of increased concern when studying developmental samples, we generated subject specific functional regions of interest (ss-fROIs) using the Group constrained Subject-Specific (GSS) method (Fedorenko et al., 2010; https://web.mit.edu/bcs/nklab/GSS.shtml). Defining ss-fROIs enhances power, and therefore, increases the chances of detecting age- and performance-related changes in MD network organization. Because motion is a concern in child samples, we asked a subset of adult subjects to participate in additional “wiggly” scans of the SWM task and to move slightly during those scans. We predicted that i) children would exhibit a similar but immature pattern of MD network activation as adults (even after accounting for motion), ii) this network would be distinct from the adjacent language network in both functional responses and connectivity, and iii) that this pattern would look more adult-like in children with better task performance regardless of their age.

## METHODS

### Participants

We recruited 44 typical adults (68% female, mean age = 23.9 years, range = 17.9-38.2 years, standard deviation = 4.9 years) and 37 typically developing children (35% female, mean age = 8.3 years, range = 4.5-12.0 years, standard deviation = 2.1 years) who completed a battery of fMRI localizer tasks as part of multiple ongoing studies investigating brain development at The Ohio State University (OSU). A total of 84 child and 50 adult scans were collected. Children who did not complete the spatial working memory (SWM) task or had data processing problems (n = 16) and those who only completed one run of the SWM task (n = 12) were not included in analyses. Seven children and one adult were excluded for excessive head motion (>25% of timepoints with >1mm total vector motion (TVM) or total framewise displacement >3 standard deviations from the group mean; Power et al., 2012). Three adults had data processing problems and two were excluded due to low accuracy (>3 standard deviations below the group mean). Of the 49 remaining child scans, we collected 2-3 scans from nine of the children, leaving a final sample of 37 children for our cross-sectional analyses. We also collected additional “wiggly” runs of the SWM task for 16 of the adults, in attempt to mimic motion in child scans, so that we could compare motion matched groups. Demographics are shown in **Table 1** for the full child and adult samples, as well as for the “wiggly” adult subgroup.

**Table 1.** Demographics and behavioral variables for kids and adults included in selectivity analyses.

|  | Adults<br>Mean(SD) or % | Wiggly Adults<br>Mean(SD) or % | Children<br>Mean(SD) or % |
| --- | --- | --- | --- |
|  | n = 44 | n = 16 | n = 37 |
| Age | 23.9(4.9) | 22.9(4.8) | 8.3(2.1) |
| Sex (% female) | 48% | 69% | 35% |
| Race |  |  |  |
| White | 71% | 65% | 74% |
| Black | 4% | 6% | 14% |
| Asian | 11% | 14% | 3% |
| Other | 14% | 15% | 9% |
| Handedness<br>(% right) | 93% | 100% | 89% |
| Framewise<br>displacement | 49.3(12.6) | 171.9(131.8) | 167.3(77.4) |

A subset of 34 children and 32 adults who also completed the language localizer task were included in analyses evaluating functional and connectivity differences between the MD and language networks. Behavioral data during the SWM task are missing or incomplete for three child and nine adult subjects, thus analyses assessing accuracy and reaction time only include 34 children and 34 adults. We also present preliminary longitudinal findings the sample of children who completed multiple scans about a year apart (n = 14; 75% female, mean age = 6.99 years, range = 4.7-9.1 years, standard deviation = 1.3 years). Five of these children do not meet our predefined motion cutoffs for at least two scans and three do not have available behavioral data during the SWM task for at least two scans. Therefore, only a subset of children scanned more than once is included in longitudinal analyses assessing change in performance (n = 6) and longitudinal analyses assessing change in resting-state connectivity (n = 9). In an interest to report findings including all children scanned more than once, results including all 14 children, regardless of motion or behavioral data missingness, are presented in supplementary materials.

Participants were recruited from the local community around OSU. Participants reported normal vision and no neurological, neuropsychologic, or developmental diagnoses at the time of recruitment. Informed consent was obtained from all participants and parental permission and assent was also obtained for all child participants. All study protocols were approved by the Institutional Review Board at OSU.

### Experimental Design

A series of functional localizers were completed by each participant, including a spatial working memory and language localizer task in the fMRI. If participants completed the full assessment, six localizer tasks (including the two tasks we report here), resting-state, T1-, and T2-weighted images were collected during an approximately 90-minute scan protocol. Parents completed a battery of parent-report measures about their child and adult participants completed self-report questionnaires.

#### Functional Localizer Tasks

A spatial working memory (SWM) task was used to functionally locate regions that respond to cognitive load, which are associated with the domain-general multiple demand network (Fedorenko et al., 2013; https://evlab.mit.edu/funcloc/). Task difficulty was adjusted by age (low, medium, and high load); all adults completed the high load version of the task which is depicted in **Supplementary Figure 1**. We contrast Hard versus Easy blocks to isolate activation selective for cognitive load. Participants viewed a grid of nine (low load) or 12 (medium/high load) boxes. Some of the boxes in the grid are blue, and participants must try to remember which boxes were blue. Participants then indicate if they think the prompt matches what they saw. Each child completed at least one run of this task outside of the scanner, to ensure they understood the task rules.

A language localizer task was used to functionally locate regions responsive to lexical and structural properties of language (Fedorenko et al., 2010; https://evlab.mit.edu/funcloc/). Participants listened to blocks of Sentences (Sn), Nonsense sentences (Ns; string of words controlling for prosody but constructed from phonemically intact nonsense words), and Texturized speech (Tx; controlling for low-level auditory features). Each run consisted of four blocks of each condition and three 14 second fixation blocks. Each block contained three trials (6 seconds each) followed by a visual queue to press a button.

### Data acquisition

Images were acquired on a Siemens Prisma 3T scanner with a 32-channel phase array receiver head coil. Foam padding used for head stabilization and increased comfort for all participants. Anatomical images were acquired with a whole-head, high resolution T1-weighted magnetization-prepared rapid acquisition with gradient echo (MPRAGE) scan to facilitate registration of masks to each subject’s anatomical space (repetition time (TR) = 1390ms; echo time (TE) = 4.6ms; voxel resolution = 1×1×1mm^3^; flip angle = 12°). Functional images during the SWM task were acquired with similar echo-planar imaging (EPI) sequences for all difficulty levels (TR = 1000ms; TE = 28ms; voxel resolution = 2×2×3mm^3^; flip angle = 61°), but the number of frames differed to provide children with longer response windows (low difficulty: 336; medium: 384; high: 400). The language localizer was acquired with the same EPI sequence for both children and adults (TR = 1000ms; TE = 28ms; Voxel resolution = 2×2×3mm^3^; number of frames = 258; flip angle = 61°). Resting-state functional images were acquired with the same I sequence for both children and adults (TR = 1000ms; TE = 0ms; voxel resolution = 2×2×3mm^3^; flip angle = 0°). Participants were asked to lay still and look at a white fixation cross on a black screen during resting-state image acquisition.

### fMRI preprocessing

#### Anatomical

Data were analyzed with Freesurfer v.6.0.0, FsFast, FSL, and custom matlab code. A semi-automated processing stream (recon-all from Freesurfer) was used for structural MRI data processing. All structural images were preprocessed using a semiautomated processing stream with default parameters (recon-all function in Freesurfer: https://surfer.nmr.mgh.harvard.edu/fswiki/recon-all/), intensity correction, skull strip, surface co-registration, spatial smoothing, white matter and subcortical segmentation, and cortical parcellation. Cortical grey matter masks for the regions of the MD network were based on prior literature (Fedorenko et al, 2013) and registered to each subject’s native anatomical space.

#### Task-based fMRI

Functional images were motion corrected (aligned all timepoints to the first timepoint in the scan and removed all time points with >1mm total vector motion between consecutive timepoints). We used bbrregister to register functional data to the subject’s anatomical space and re-sampled to 1×1×1mm^3^. Volume-based, first-level general linear model analyses applied additional processing: detrended and smoothed (SWM task: 4mm FWHM kernel; Language localizer: 5mm FWHM kernel). A standard boxcar function was used to (events on/off) convolve the canonical hemodynamic response function (standard gamma function (d=2.25 and t=1.25)) and included a regressor for each condition (SWM task: Hard, Easy; Language localizer: Sn, Ns, Tx). Additional nuisance regressors (six orthogonalized motion measures from the preprocessing stage) were applied to the processed images for each task individually. Resulting beta estimates and contrast maps (SWM task: Hard > Easy; Language localizer: Sn > Ns) were used for subsequent analyses.

#### Resting-state fMRI

We preprocessed the resting-state data using Freesurfer’s FS-Fast preprocessing pipeline (https://surfer.nmr.mgh.harvard.edu/fswiki/FsFastAnlysisBySteps). Framewise displacement was used as a motion regressor. We generated masks of white matter, cerebrospinal fluid (CSF), and subcortical structures for each subject in their native anatomical space. We performed spatial smoothing, interpolation over motion spikes, bandpass filtering (0.009-0.08 Hz), and denoising using CSF and white matter masks. Timepoints with framewise displacement greater than 0.5mm were censored from the data.

### Generating a probabilistic map

We smoothed each subject’s significance map (4mm FWHM kernel) and registered to FsAverage space (https://surfer.nmr.mgh.harvard.edu/fswiki/FsAverage). We binarized and thresholded each subject’s significance map (-log10(p) > 2; or p <0.01) and summed the number of subjects who exhibit significant activation in each voxel.

### Subject-specific functional regions of interest (fROI)

We used the Group constrained Subject-Specific method (GSS; Fedorenko et al., 2010) to define subject-specific functional regions of interest (ss-fROIs). Fedorenko and colleagues (2013) derived 20 regions (10 bilateral regions) belonging to the MD network based on probabilistic maps of functional activation in a sample of adults (i.e., grey matter masks). Each region included showed greater functional activation during Hard compared to Easy blocks in at least 60% of their subjects across six different cognitively demanding tasks. Each of the masks were registered using Freesurfer’s CVS function (Postelnicu, Zollei, & Fischl, 2008; https://surfer.nmr.mgh.harvard.edu/fswiki/mri_cvs_register) from the CVS atlas MNI152 space to the participant’s native anatomical space. We used one run of the first level GLM analysis to identify each subject’s ss-fROIs by assessing subject-specific contrast maps within all 10 bilateral search spaces. The top 10% of activated voxels during the contrast of interest (i.e., Hard > Easy) were considered the ss-fROI. Two sets of ss-fROIs were generated for each subject (i.e., one for each run of the SWM task).

### Percent signal change (PSC)

#### Cross-sectional

Percent signal change (PSC) for each condition (SWM task: Hard, Easy; Language localizer: Sn, Ns, Tx) was extracted within all ss-fROIs generated from an independent run of the SWM task (e.g., PSC during run 1 was extracted within the ss-fROIs generated from run 2) using the first-level beta estimates. Ss-fROIs generated from the SWM task were used to extract PSC from the Language localizer task for the 35 children and 32 adults who also completed at least one run of the Language localizer task (and PSCs were averaged across runs if they completed more than one run).

#### Longitudinal

For the 14 subjects who completed two longitudinal scans, we registered ss-fROIs defined at their second timepoint to the subject’s native anatomical space at their first timepoint using Freesurfer’s CVS function (Postelnicu, Zollei, & Fischl, 2008). We then used these timepoint two ss-fROIs (that were now in timepoint one’s native anatomical space) to extract PSC during the SWM task from timepoint one so that change in selectivity for cognitive load could be calculated in the same ss-fROIs over time. For subjects with two runs of the SWM task at timepoint one, the ss-fROI generated from the first run at timepoint two was used to extract PSC from the first run at timepoint one; this was repeated for the second run of the SWM task and the PSC values were averaged. For subjects with only one run of the SWM task at timepoint one, the ss-fROI generated from the first and second runs at timepoint two were used to extract PSC from the first run at timepoint one and then averaged.

### Selectivity Indexes

The MD selectivity index quantifies selectivity of MD ss-fROIs for cognitive demand (**Formula 1**). Reported MD selectivity indexes reflect averaged selectivity between the first and second runs of the SWM task. The Language selectivity quantifies selectivity of MD ss-fROIs for language (**Formula 2**). Reported language selectivity indexes reflect selectivity during the first run for those who only completed one run of this task (n = 7) and averaged selectivity between the first and second runs of the Language localizer task for participants who completed two runs (n = 27).

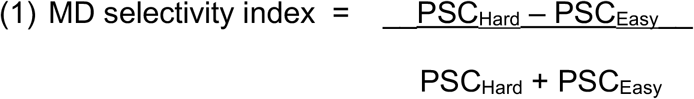

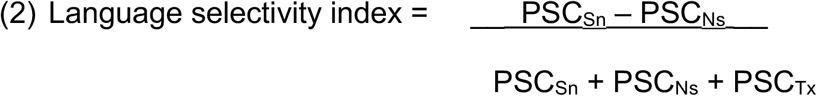

### Resting state connectivity

We generated ss-fROIs for each subject that completed the SWM task and Language localizer task. The average time course for each ss-fROI of the MD and language networks was computed from the pre-processed resting-state images. To evaluate interactions between regions, Pearson correlations were generated between ss-fROI time courses. To generate normally distributed values, each functional connectivity value was Fisher z-transformed. For the 35 children that completed both tasks, we compared mean within- and between-network connectivity using repeated measures t-tests.

### Statistical analyses

Single sample sign tests were conducted to evaluate if selectivity is greater than zero because of the positively skewed distribution of MD selectivity within the child sample. We report the dominance statistic (DS) as a measure of effect size for MD selectivity values, which reflects the proportion of the sample exhibiting MD selectivity greater than zero minus the proportion exhibiting MD selectivity less than zero (Mangiafico, 2016). Independent-sample t-tests (due to unequal sample sizes) were conducted to evaluate whether MD selectivity differs between the motion matched adults and children. Paired sample sign tests were conducted to evaluate MD selectivity versus language selectivity in MD ss-fROIs for children, MD selectivity during wiggly vs. non-wiggly runs for a subgroup of adults, and within-network versus between-network (with language network ss-fROIs) resting-state connectivity. We corrected for multiple comparisons using Bonferroni-Holm correction; we corrected for three frontal and seven parietal comparisons in each hemisphere.

To evaluate linear relationships between selectivity, age, and task performance we conducted Pearson’s correlations and multiple linear regressions for the cross-sectional analyses and conducted linear mixed effects models (to account for within-subject shared variance) for the longitudinal analyses (lmerTest R package; Luke, 2017). All selectivity analyses were conducted in RStudio Version 1.1.331 and resting-state statistical analyses were performed in MATLAB Version R2020a.

### Code Accessibility

We used custom scripts in MATLAB for much of the data processing (e.g., generating ss-fROIs and extracting PSC). Scripts are available upon request; contact corresponding author (ZS).

## RESULTS

### Exploratory probabilistic map of the multiple demand network in children

We generated an exploratory probabilistic map of voxels that exhibit significantly greater activation (z > 2.58) during Hard versus Easy trials of a spatial working memory task for at least two subjects (**Figure 1**). The maximum number of subjects who show significant activation in the same voxel is eight (22% of cross-sectional sample). Despite functional variability, we see a pattern of activation across several frontal and parietal regions that is similar to the pattern seen in adults (Fedorenko et al., 2013).

**Figure 1.**
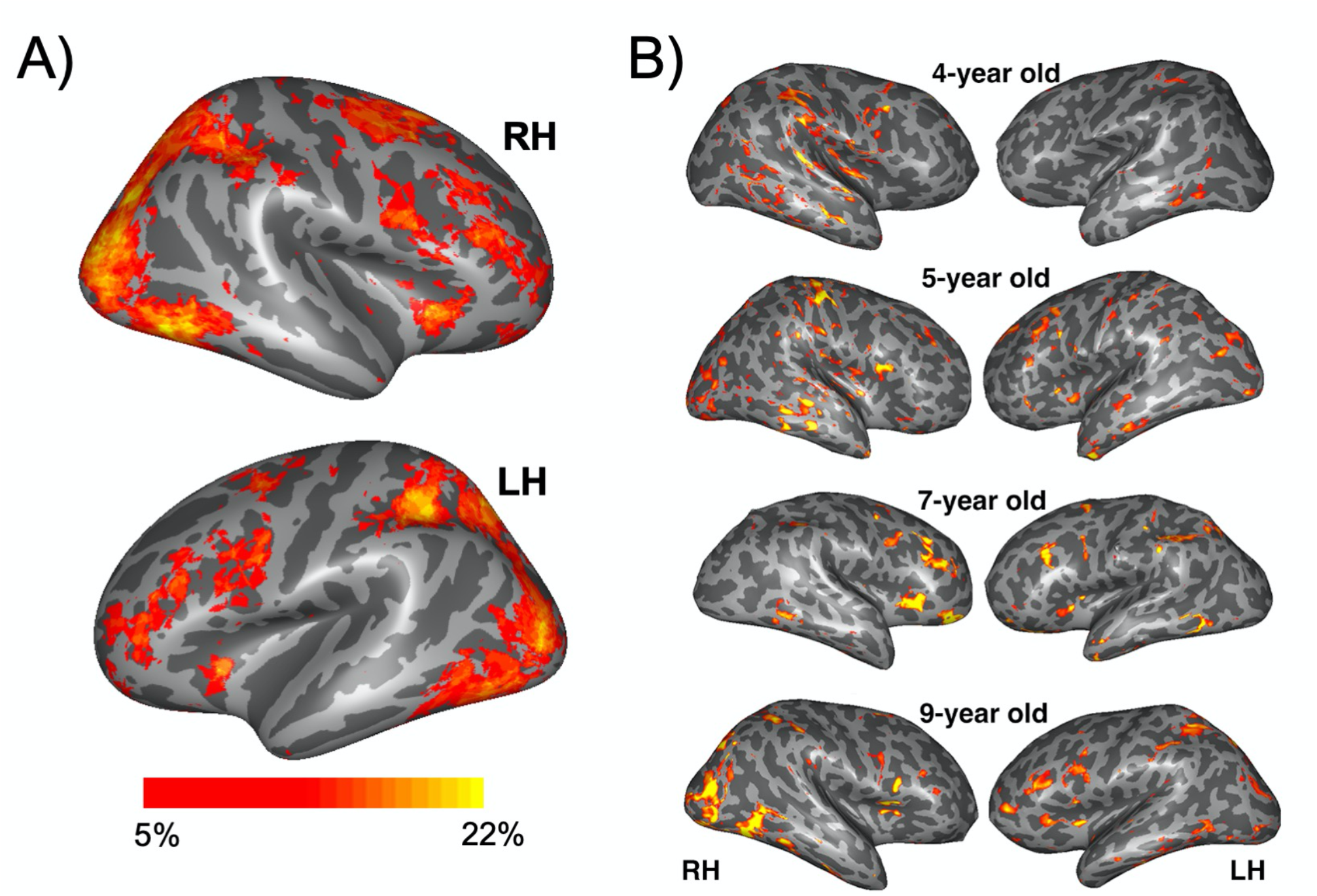
A) Probabilistic map showing common activation during a blockwise spatial working memory task (z > 2.58; contrast: Hard–Easy) in at least two participants and a maximum of eight participants (22% of participants). B) Example significant maps are depicted for four participants.

### Selectivity of the multiple demand network in adults

We considered ss-fROIs to be selective to cognitive demand if their neural responses to Hard trials (on a spatial working memory task) was higher than to Easy trials (i.e., MD selectivity index was significantly greater than zero). We tested this using single sample, one-tailed, sign tests. **Table 2** shows group descriptive statistics and p-values for each MD region’s MD selectivity index and Figure 2 provides a visual depiction of mean selectivity by region and group. As expected, adults show selectivity in all bilateral MD ss-fROIs (uncorrected *p* < .001; Bonferroni-Holm corrected *p* < .05 for all ss-fROIs; **Table 2**). However, motion is a problem in child neuroimaging, which can cause spurious correlations (Power et al., 2012). Even with strict cut-offs like our current sample (see Methods), children may still move more than adults. We asked a subset of our adult participants to complete additional runs of the SWM experiment while encouraging them to move in the scanner (‘wiggly adult runs’). This group was not significantly different in total framewise displacement from the child group (**Table 1**). Further, only the left precentral gyrus shows significantly greater selectivity in wiggly versus non-wiggly runs within the adult sample (*p* = .021, 95% CI for difference in scores [-0.49, -0.07], repeated measures sign test), which did not survive multiple comparison correction (descriptive statistics shown in **Table 2**). In this motion-matched wiggly adult group, all bilateral MD ss-fROIs remain significant (uncorrected *p* < .05); all bilateral parietal, all right frontal, and half of the left frontal MD ss-fROIs survive multiple comparison correction (Bonferroni-Holm corrected *p* < .05). This suggests that selectivity of MD ss-fROIs is robust among adults, even in the face of child-like motion and suggests that any developmental differences we observe are likely not driven by motion.

**Table 2.**
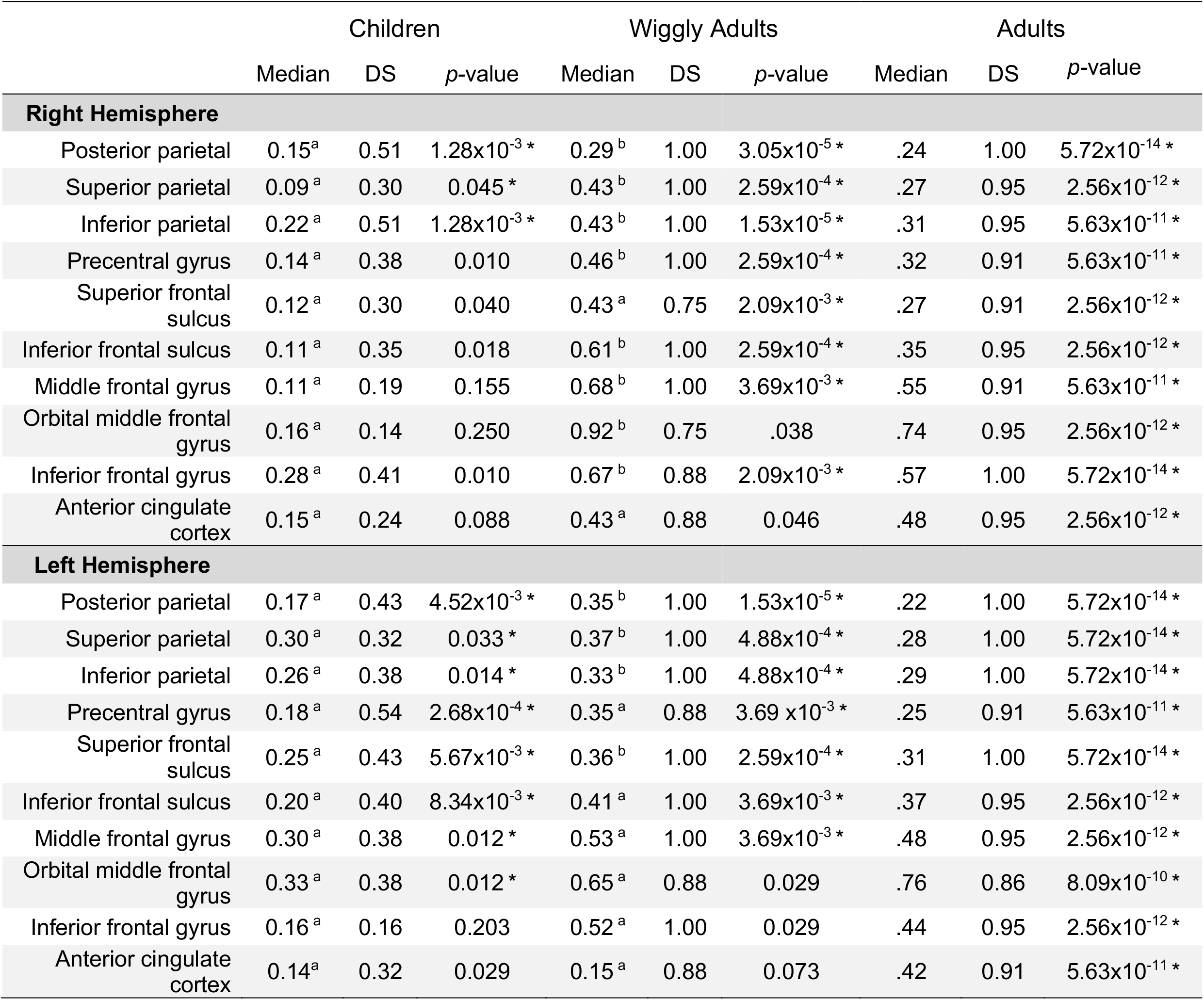
Selectivity of the multiple demand (MD) ss-fROIs during a spatial working memory task. One-tailed sign-tests were conducted and medians are reported due to positively skewed distribution. Regions are grouped by hemisphere and brain lobe. Asterisk indicates corrected *p* < .05 (Bonferroni-Holm, 3 parietal regions, 7 frontal regions). ^a, b^ notations indicate significant differences between the motion matched child and adult groups (uncorrected *p* < .05, two-tailed t-tests). DS = Dominance Statistic.

**Figure 2.**
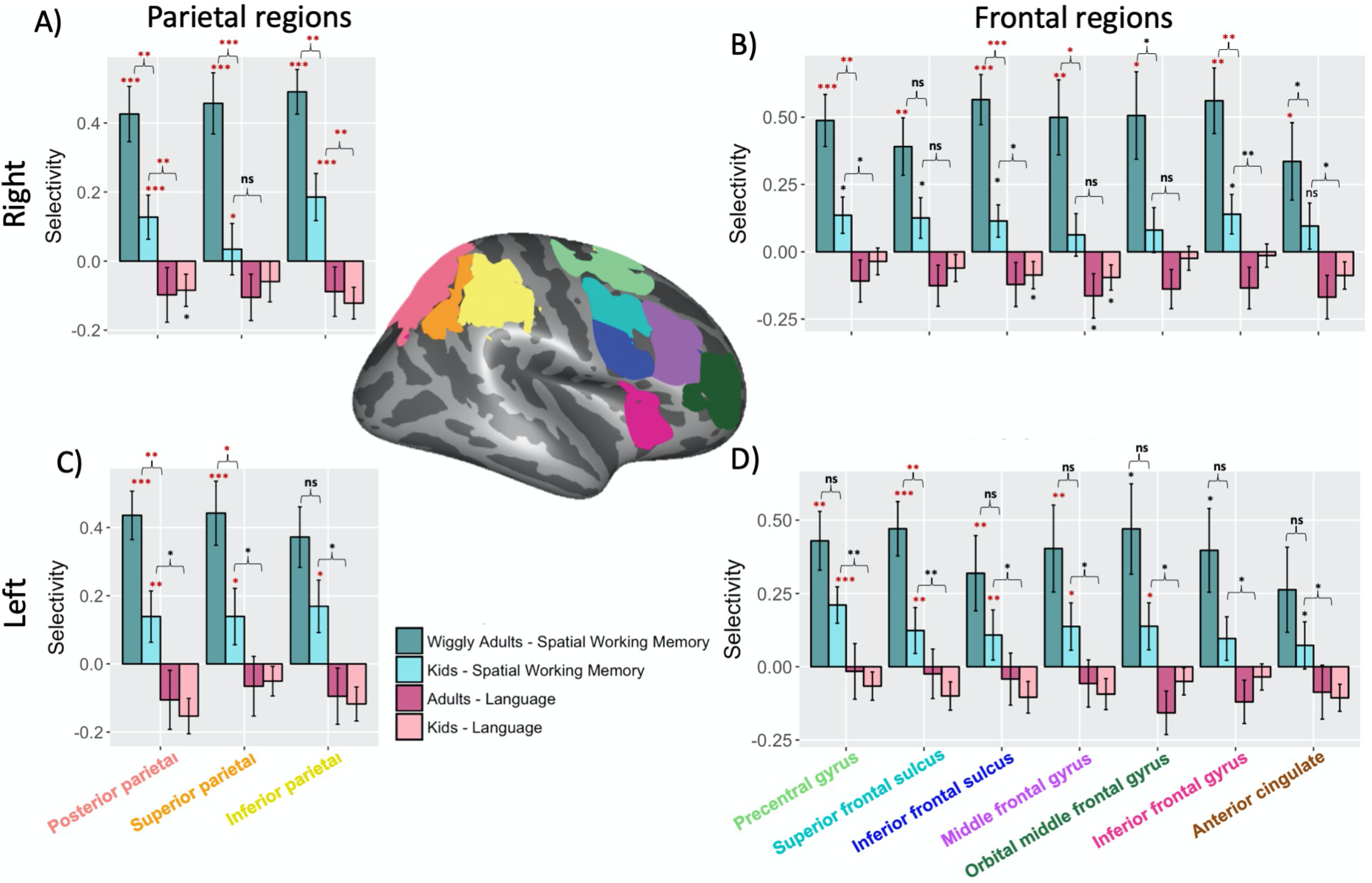
Mean selectivity in the right (A,B) and left (C,D) multiple demand (MD) network ss-fROIs during the spatial working memory (blue) and language localizer (pink) tasks for children (lighter colors) and adults (darker colors). One-tailed sign tests were conducted to test if MD selectivity is significantly greater than zero, two-tailed sign tests were conducted to test if language selectivity is significantly different from zero, repeated measures sign tests were conducted to test if children show greater MD than language selectivity in MD ss-fROIs, and independent t-tests (due to uneven group sizes) were conducted to test if adults show greater MD selectivity than children in motion matched groups. Asterisks indicate strength of significance (uncorrected): * *p* < .05; ** *p* < .01; *** *p* < .001. Red symbols indicate Bonferroni-Holm corrected p < .05, corrected for 10 comparisons per hemisphere. Error bars denote standard error.

### Selectivity of the multiple demand network in children

A similar pattern is observed among children. Most MD ss-fROIs show positive selectivity indexes (**Figure 2**; light blue bars). Children exhibit significant selectivity in all three bilateral parietal, four right frontal, and six left frontal MD ss-fROIs (uncorrected *p* < .05; **Table 2**). The bilateral posterior parietal, inferior parietal, and superior parietal and the left precentral gyrus, inferior/superior frontal sulci, middle frontal gyrus, and orbital middle frontal gyrus survive multiple comparison correction (Bonferroni-Holm corrected *p* < .05). Only three right (anterior cingulate [though trending], orbital middle frontal gyrus, middle frontal gyrus) and one left (inferior frontal gyrus) frontal regions do not reach significance. These data show that in samples of children as young as four years we can already see functionally specialized activation of the MD network during a cognitive demanding task.

How does this selectivity compare to adults? Upon visual inspection, all ss-fROIs show greater selectivity in adults as compared to children (**Figure 2**), but surprisingly only 11 of the 20 ss-fROIs reach statistical significance for (uncorrected *p* < 0.05, independent sample, two-tailed, t-tests; **Table 2**). Eight MD ROIs in the right hemisphere (all three parietal, five frontal) show significantly greater selectivity in adults compared to children (uncorrected *p* < .05); seven of these ss-fROIs (right inferior frontal sulcus, precentral gyrus, inferior frontal gyrus, middle frontal gyrus, superior parietal, inferior parietal, posterior parietal) survive multiple comparison correction (Bonferroni-Holm corrected *p* < .05; **Figure 2**). On the contrary, only one frontal and two parietal ss-fROIs in the left hemisphere are significantly greater in adults (uncorrected *p* < .05) and only three (superior frontal sulcus, posterior parietal, superior parietal) survive multiple comparison correction. Thus, while adults evoke stronger selectivity than children, the selectivity differences are not extensive and widespread, and these MD ss-fROIs still show strong selectivity in young children.

These data show that even in young children, the bilateral MD network is engaged during demanding tasks, just like in adults. Though we see more activation during Hard trials of a spatial working memory task than Easy trials in both children and adults, we still see greater selectivity in adults than in children, primarily among right hemisphere ss-fROIs. These differences do not appear related to motion and may reflect maturation of these ss-fROIs.

### Dissociation between MD and adjacent language regions: functional selectivity

We next focused more closely on the frontal ss-fROIs and compared the same participants’ activation on this task to activation on a language task, where participants were asked to listen to meaningful Sentences vs. Nonsense sentences. Frontal MD and language regions are often in close proximity, but we know from previous work (Fedorenko et al., 2013) that these networks are functionally distinct in adults. Do these regions emerge from a common frontal area? If so, we may expect slight sensitivity, or mild selectivity, to language in these MD ss-fROIs in young children.

We conducted single sample, two-tailed, sign tests to test this hypothesis. We found that no MD ss-fROIs show selectivity to language in either our child or adult samples (i.e., no ss-fROIs were significantly greater than zero). Intriguingly, we see trends of greater signal during the Nonsense sentences condition than the Sentences condition (reflected as language selectivity less than zero in the contrast of English>Nonsense; **Figure 2**, pink bars). Several ss-fROIs (bilateral inferior frontal sulcus and posterior parietal, right middle frontal gyrus and anterior cingulate, and left inferior parietal) reach significance in our child sample and one region in our adult sample (left middle frontal gyrus; uncorrected *p* < 0.05). However, none survive multiple comparison correction. Nonsense sentences may evoke greater functional activation in regions associated with cognitive demand than language regions as individuals attempt to unscramble the words, effectively engaging in a cognitively demanding task, instead of language processing.

We also asked whether children show greater MD selectivity than language selectivity in these ss-fROIs using repeated measures, one-tailed, sign tests. All left hemisphere ss-fROIs show greater MD than language selectivity, and half of the right hemisphere MD ss-fROIs reach significance for this comparison (uncorrected *p* < .05); the right posterior and inferior parietal MD ss-fROIs survive multiple comparison correction (Bonferroni-Holm corrected *p* < .05). Overall, we see selective activation for spatial working memory in the MD network but no language selectivity within these ss-fROIs in children. Are these networks dissociated in their connectivity patterns as well?

### Dissociation between MD and adjacent language regions: connectivity

Previous work shows that in adults, regions of the multiple demand and language networks are also distinct in resting-state functional connectivity (Blank, Kanwisher, & Fedorenko, 2014). We investigated whether this pattern is also observed among children using the MD ss-fROIs defined above and also defined separate language ss-fROIs using the language task in each child. We find significantly greater within-MD than between-network connectivity (i.e., between regions of the MD and language networks) in both the left (t(33) = 3.37, p = 1.90×10^−3^) and right (t(33) = 3.27, p =2.5×10^−3^) hemisphere (**Figure 3A**). Focusing on just the frontal ss-fROIs (which exist in both networks), we also find the same pattern: frontal MD ss-fROIs are more connected to one another than they are to adjacent frontal language ss-fROIs in both the left (t(33) = 3.46, p = 1.90×10^−3^, repeated measures t-test) and right (t(33) = 2.87, p = 7.1×10^−3^, repeated measures t-test) hemispheres. We see a similar pattern in our longitudinal sample (n = 9). At timepoint one, children show marginally greater within-than between-network connectivity in both hemispheres (left: t(8) = 1.93, p = .090; right: t(8) = 2.16, p = .063, repeated measures t-tests; **Figure 3B**) and at timepoint two within-network connectivity is significantly greater than between-network connectivity in both hemispheres (left: t(8) = 4.10, p = 3.4×10^−3^; right: t(8) = 2.34, p = .047, repeated measures t-tests; **Figure 3C**). These findings are consistent with our hypothesis that the MD network is already dissociated in both function and connectivity from the adjacent language network in young children.

**Figure 3.**
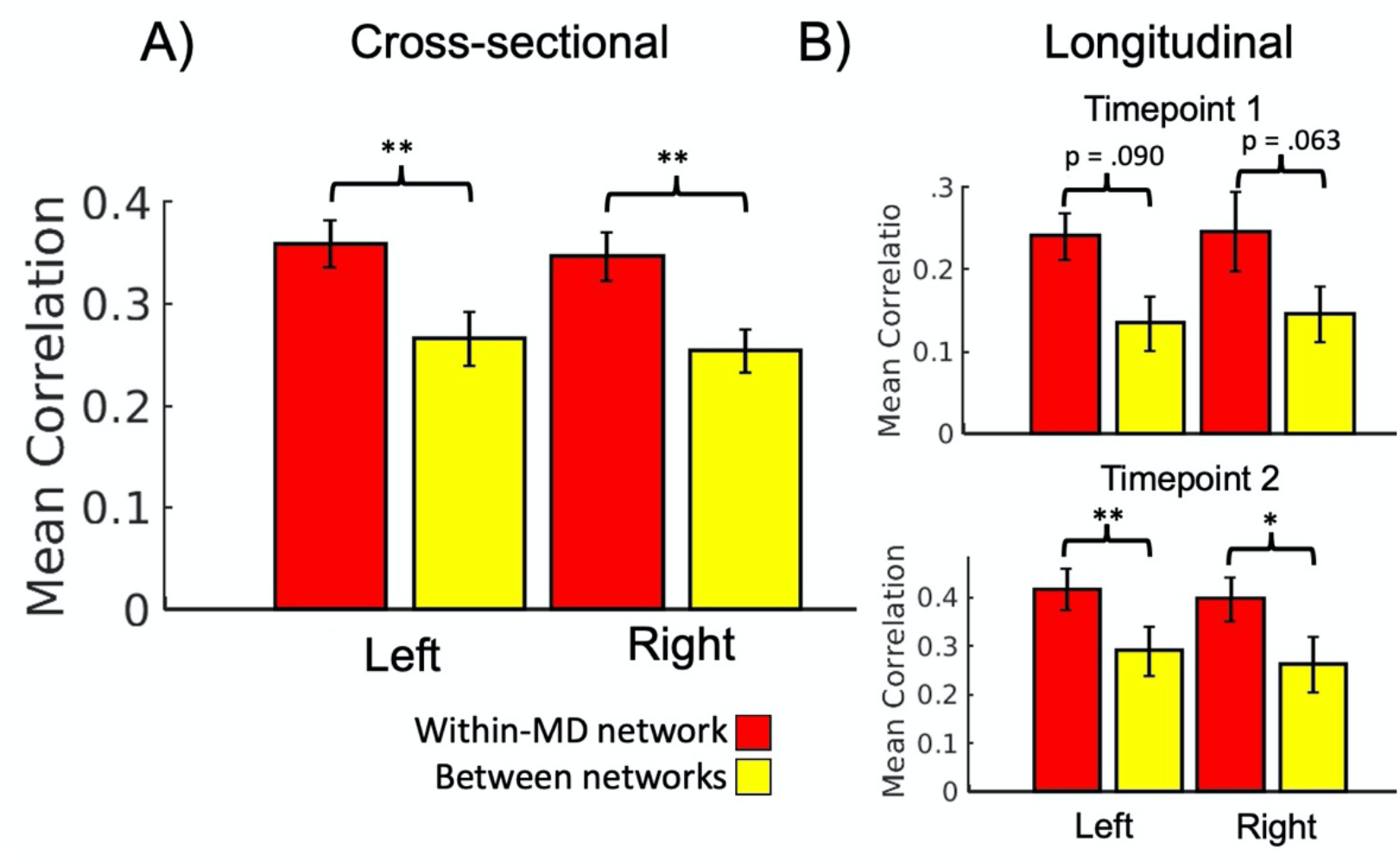
Mean resting-state connectivity between subject-specific function regions of interest (ss-fROIs) for the multiple demand (MD) and language networks, separated by hemisphere, the cross-sectional (A) and longitudinal (B) samples. Repeated measures t-tests show significantly greater within-than between-network connectivity in both hemispheres. Asterisks indicate strength of significance (uncorrected): * p < .05; ** p < .01. Error bars denote standard error.

We then assessed whether connectivity is associated with age. In the cross-sectional sample we do not see a correlation with age of within-MD network connectivity in either hemisphere (*p* > .14, Pearson’s correlations). In the longitudinal sample, we see a significant correlation between age and within-MD network connectivity in the right hemisphere (within-all MD: B = 0.11, t(13.7) = 3.91, p =1.65×10^−3^; within-frontal MD: B = 0.10, t(15.7) = 2.56, p =.021, linear mixed effects models), but not the left hemisphere (p >.19). There were no significant increases with age in within minus between-network connectivity for either hemisphere in the cross-sectional (p > .50, Pearson’s correlation) or longitudinal sample (p > .39, linear mixed effects model). This suggests that the MD network is mostly dissociated from the language network already during early-to-mid-childhood.

To assess how within-MD network connectivity is related to current ability, regardless of age, we assess the association between within-MD network connectivity and accuracy during the SWM task while control for age. After controlling for variance explained by age, the models show no association between within-MD network connectivity and accuracy in either hemisphere for the cross-sectional (p > .29, multiple linear regression models) and longitudinal (p > .38, linear mixed effects models) samples. These data suggest that connectivity of the MD network is more related to age, than to EF performance. To explore age- and performance-related associations further, we examine task-dependent functional selectivity of MD ss-fROIs during the SWM task.

### MD selectivity is not associated with age

Despite similarities to adults, differences do exist between children and adults, such as in the selectivity of the MD ss-fROIs (**Figure 2**). Does the MD network change in selectivity with age in our sample of children from 4-12 years? Or does it reflect individual differences in performance, regardless of age? We explored this question both cross-sectionally and longitudinally in our sample. We first investigated whether selectivity was associated with age in our cross-sectional sample. Interestingly, MD selectivity was not significantly associated with age for the majority of MD ss-fROIs (p > .06, Pearson correlation). We also performed analogous correlations in our longitudinal sample, and, similarly, we observed no significant associations with age in the model with age alone (p > .09, linear mixed effects models).

### MD selectivity is associated with performance

We then asked whether selectivity is more related to individual variability in performance, rather than age. During Hard trials in the cross-sectional sample, we found that one bilateral parietal, one right parietal, one left parietal, one left frontal, and five right frontal ss-fROIs show increasing MD selectivity with performance (uncorrected p < .05, Pearson correlations; **Supplemental Table 1**). Five of the right ss-fROIs (middle frontal gyrus, inferior/superior frontal sulcus, inferior parietal, and precentral gyrus) and two left parietal ss-fROIs (posterior/superior parietal) survive multiple comparison correction (Bonferroni-Holm corrected p < .05; three ss-fROIs’ correlations are depicted in **Figure 4**). We see a similar pattern of associations for MD selectivity and Easy trial performance (**Supplemental Table 1**).

**Figure 4.**
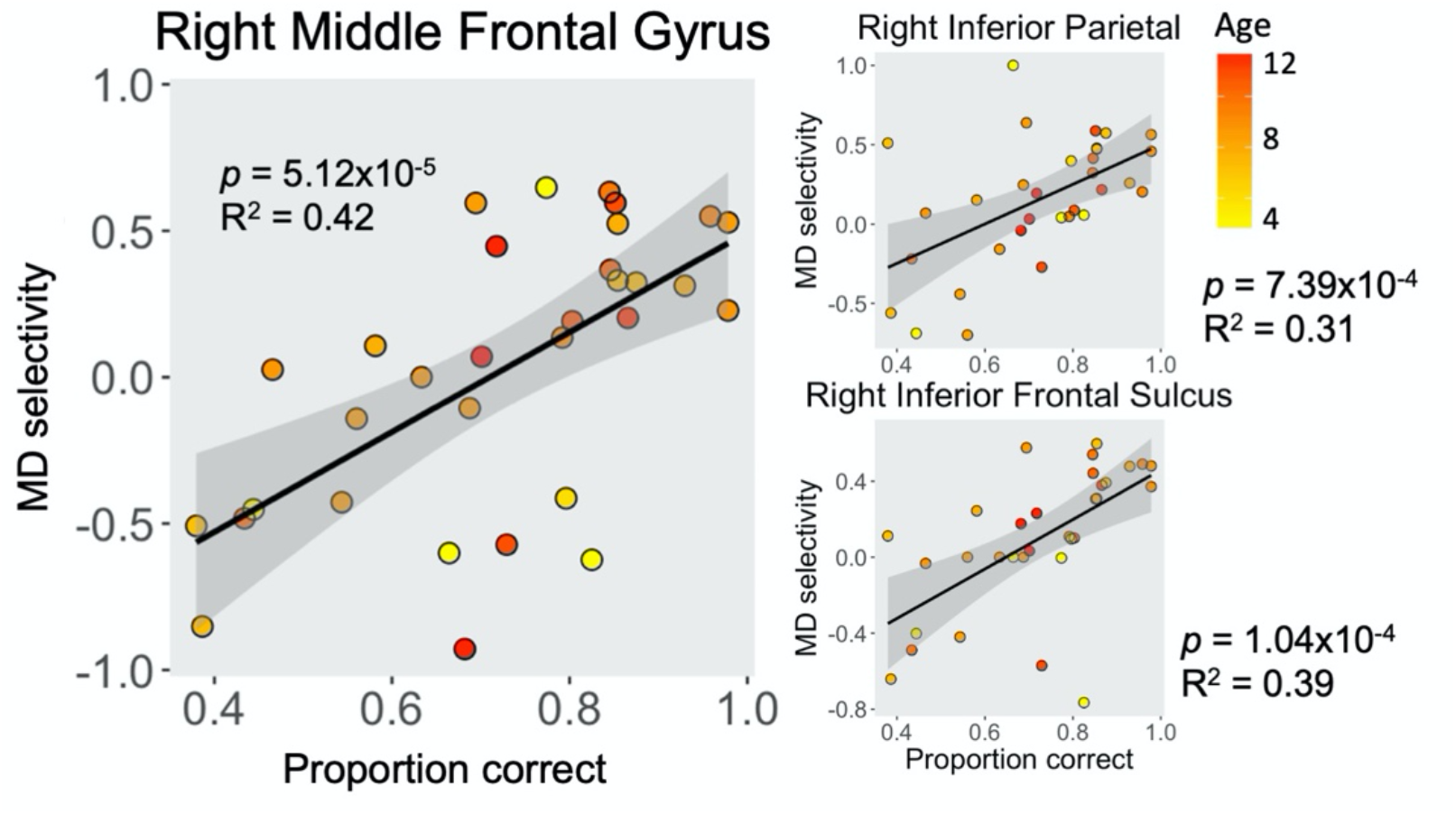
Positive association between neural selectivity of MD network and behavioral accuracy during Hard trials of the SWM task; stronger selectivity was related to better task performance regardless of age. Bonferroni-Holm corrected *p* < .05 for all three ss-fROIs shown here; two left ss-fROIs also survive multiple comparison correction.

To ensure that these results are not confounded by age, especially given that accuracy during Easy trials is positively related to age (r = 0.44, t(30) = 2.67, p = .012, Pearson correlation; **Supplemental Figure 2**), we controlled for age in subsequent models. We did not control for framewise displacement because no ss-fROI MD selectivity showed significant correlation with framewise displacement in the cross-sectional sample (p > .09). During Hard trials, seven right ss-fROIs and two left ss-fROIs remain significantly related to accuracy after controlling for age (**Table 3**); three of the right ss-fROIs (inferior frontal sulcus, middle frontal gyrus, inferior parietal) survive multiple comparison correction (Bonferroni-Holm corrected p < .05). Only the right posterior parietal and anterior cingulate (**Table 3**) show a significant main effect of age for significant models including accuracy (Hard trials) and age. We see a similar pattern of associations between selectivity and accuracy during Easy trials (**Supplemental Table 2**).

**Table 3.** Main effect of accuracy during Hard trials on selectivity of MD ss-fROIs during a spatial working memory task for our child sample. Multiple linear regressions were conducted to account for variability associated with age. Asterisks indicate strength of significance (uncorrected): * p < .05; ** p < .01; *** p < .001; ^m^ Bonferroni-Holm full model corrected p < .05 (3 parietal regions, 7 frontal regions).

| | Accuracy<br>B | Accuracy<br>t-value | Age<br>B | Age<br>t-value | Full model<br>F-value | Adjusted<br>$R^2$ |
| --- | --- | --- | --- | --- | --- | --- |
| **Right Hemisphere** |  |  |  |  |  |  |
| Posterior parietal | 0.69 | 1.98 | -0.06 | 2.22* | 3.83* | 0.15 |
| Superior parietal | 1.00 | 2.10* | -0.009 | -0.23 | 2.20 | 0.07 |
| Inferior parietal | 1.30 | 3.63** | -0.02 | -0.85 | 6.64** | 0.27<sup>m</sup> |
| Precentral gyrus | 1.18 | 3.10** | 0.008 | 0.27 | 5.07* | 0.21 |
| Superior frontal sulcus | 1.19 | 2.64* | 0.008 | 0.23 | 3.69 | 0.15 |
| Inferior frontal sulcus | 1.25 | 4.01*** | 0.03 | 1.12 | 9.56*** | 0.36<sup>m</sup> |
| Middle frontal gyrus | 1.66 | 4.21*** | 0.02 | 0.76 | 9.88*** | 0.36<sup>m</sup> |
| Orbital middle frontal gyrus | 0.83 | 1.62 | 0.06 | 1.43 | 2.75 | 0.10 |
| Inferior frontal gyrus | 0.46 | 1.06 | 0.02 | 0.67 | 0.91 | -0.01 |
| Anterior cingulate cortex | 0.92 | 1.98 | 0.09 | 2.23* | 5.23* | 0.21 |
| **Left Hemisphere** |  |  |  |  |  |  |
| Posterior parietal | 1.11 | 2.87** | -0.04 | -1.37 | 4.58* | 0.19 |
| Superior parietal | 1.19 | 2.38* | -0.007 | -0.16 | 2.85 | 0.11 |
| Inferior parietal | 0.86 | 1.74 | 0.03 | 0.65 | 1.95 | 0.06 |
| Precentral gyrus | 0.40 | 1.07 | -0.01 | -0.44 | 0.61 | -0.03 |
| Superior frontal sulcus | 1.02 | 2.05* | 0.02 | 0.51 | 2.44 | 0.08 |
| Inferior frontal sulcus | 0.74 | 1.36 | 0.08 | 1.85 | 3.09 | 0.12 |
| Middle frontal gyrus | 0.83 | 1.61 | 0.04 | 0.90 | 1.97 | 0.06 |
| Orbital middle frontal gyrus | 0.24 | 0.47 | 0.06 | 1.36 | 1.16 | 0.01 |
| Inferior frontal gyrus | 0.41 | 0.92 | 0.08 | 2.16* | 3.14 | 0.12 |
| Anterior cingulate cortex | 0.85 | 1.76 | 0.07 | 0.07 | 3.90* | 0.16 |

Adults also show significant correlations between selectivity and accuracy during Hard trials for many of the same regions: right inferior frontal sulcus (r = .41, t(33) = 2.58, p = 0.014, Pearson correlation), right middle frontal gyrus (r = 0.34, t(33) = 2.06, p = 0.048), bilateral inferior frontal gyrus (left: r = 0.39, t(33) = 2.43, p = 0.021; right: r = 0.53, t(33) = 3.55, p = 1.18×10^−3^), and right anterior cingulate (r = 0.34, t(33) = 2.0786, p = 0.046). No significant associations were observed between selectivity and accuracy during Easy trials in the adult sample (p > .05). We did not control for age, given that accuracy in adults was not associated with age for either Hard or Easy trials (p > .31). Overall, these results suggest that children are already showing adult-like individual variability in performance which is related to neural activation on this task. The most robust associations between selectivity of MD ss-fROIs and performance (i.e., accuracy) are observed during Hard trials in right frontal regions.

Lastly, we examined these age and performance associations in our longitudinal sample. Age is significantly correlated with accuracy during Hard (B = 0.08, t(9.6) = 2.85, p = .018) and Easy (B = 0.09, t(6.8) = 4.38, p = 3.53×10^−3^) trials within this longitudinal sample (n = 6). The mixed effects models show associations between accuracy (Hard) and selectivity for some of the same regions that were significant in the cross-sectional sample, while controlling for age and framewise displacement (because a few regions showed correlations between framewise displacement and selectivity in the longitudinal sample, even after motion cutoffs were applied). We see a significant association between accuracy during Hard trials in the right middle frontal gyrus (B = 3.94, t(4.2) = 3.36, p = 2.65×10^−2^) and left posterior parietal cortex (B = 3.14, t(3.8) = 3.57, p = 2.57×10^−2^; **Supplemental Table 3**). Age and framewise displacement also show significant main effects on selectivity for some MD ss-fROIs (including the right middle frontal gyrus and left posterior parietal cortex) beyond the variance explained by accuracy (**Supplemental Table 3**). We also see a significant association between MD selectivity and accuracy during Easy trials in several MD ss-fROIs (right inferior frontal gyrus, orbital middle frontal gyrus, inferior/superior frontal sulcus, and in the left superior parietal and anterior cingulate cortex (**Supplemental Table 3**). Findings from the full sample of longitudinal children (n = 14; no motion cutoffs and some missing behavioral data) are discussed in the supplemental materials (**Supplemental Table 4**). Motion was the most robust predictor of MD selectivity in full longitudinal sample, highlighting the importance of accounting for motion when examining developmental samples and comparing them to adults. These results suggest that relationships between performance and MD selectivity replicate, even in a small sample of motion-matched longitudinal kids. This suggests that children’s neural responses reflect their EF skills and look increasingly more adult-like with increasing EF ability across time.

## DISCUSSION

Do children process cognitively demanding tasks using the same neural system as adults? Meta-analyses report similar frontoparietal regions recruited across EF components (i.e., working memory, shifting, inhibition) in both adults (Niendam et al., 2012) and children older than six years (McKenna et al., 2017), suggesting that EF is supported by a domain-general network (so-called Multiple Demand, or MD network) which is engaged across all of these EF tasks in children and adults alike. But is the same network also engaged in children as young as four years? Is this network distinct from adjacent language-specific network or are these not yet disentangled in early development? And does the recruitment of this network predict cognitive effort exerted on the task (like it does in adults) or does it instead reflect general maturation of these frontoparietal regions? These questions have not yet been explored in prior literature.

Using two fMRI tasks and resting-state fMRI we defined ss-fROIs delineated in subject-specific space (ss-fROIs) and examined task activation and connectivity in young children. By using ss-fROIs, we increased the power of our analyses by accounting for individual variability in neural structure and organization. We also analyzed both cross-sectional and longitudinal samples of children so that we could evaluate these relationships using different approaches, compare, and draw conclusions based on consistent findings between different designs and statistical tests. Further, we used strict motion-cutoffs for children and adults, and additionally acquired motion matched data in the same sample of adults, which provides more evidence that any differences observed between children and adults were not driven by motion. We found that children, like adults, recruit MD ss-fROIs while engaged in a cognitively demanding task. We observed positive selectivity (reflected by significantly greater activation for Hard vs. Easy trials) in all bilateral parietal regions and many bilateral frontal regions in our sample of children. Although many of these regions already show positive selectivity, we still see group differences between motion matched child and adult groups, possibly reflecting children’s immature EF capabilities.

Motion-matched adults show stronger selectivity in all MD regions compared to children, but not all differences were statistically significant. Interestingly, the most robust differences between children and adults were in the right hemisphere. This may speak to the commonly observed right lateralization in adults during visuospatial processing (Crittenden & Duncan, 2014; de Schotten et al., 2011; Jiang & Kanwisher, 2003). We did not observe lateralization in children and speculate that right lateralization likely develops later in childhood or adolescence. This is consistent with meta-analytic findings that right lateralization was only observed in adolescent, but not child, subgroups (Houdé et al., 2010).

Because of observed age-group differences, we also examined how functional selectivity is related to age and task performance in our child and adult samples separately. Does selectivity of MD ss-fROIs look more adult-like with increasing age, or can this variability be better explained by individual variability in task performance? Surprisingly, we found few significant associations with age. Age related changes in working memory behavioral metrics (i.e., accuracy, reaction time) are observed between children, adolescents, and young adults, such that response time and accuracy show linear negative and positive associations, respectively, with chronological age (Linares et al., 2016; Gathercole et al., 2004). We also observed similar age-related changes in accuracy and reaction time; we therefore controlled for age when assessing performance related changes in selectivity and found that selectivity is more associated with performance on the working memory task than to chronological age. Some of these significant relationships even hold in a small longitudinal sample of six children, who show significant relationship between accuracy and the right middle frontal gyrus, which seems to be the most consistent finding across samples. We also see these performance related associations in our adult sample, primarily in right frontal regions. This is consistent with findings that individual differences in variability of EF performance are related to changes in MD network activation for an independent sample of adults (Assem, Blank, Mineroff, Ademoğlu, & Fedorenko, 2020). In a sample of older children, adolescents, and young adults, evidence of a stronger relationship between working memory performance and MD network activation also exists (Satterthwaite et al., 2013). Moreover, Satterthwaite and colleagues (2013) show that MD network activation mediates the relationship between age and task performance. This suggests that functional selectivity of the MD network is a potential neural mechanism underlying age related improvement in working memory performance throughout development.

Of note, some previous studies have found age related activation changes, but not performance related changes, e.g., in a sample of 7-22-year-olds engaging in a working memory task (Kwon et al., 2002) and meta-analytic work including 4-17-year-olds engaging in various EF tasks (Houdé et al., 2010). These studies included a wider age range than our sample, thus our findings do not contradict age-related changes (independent of ability) in older samples. Because variability in performance decreases with increasing age (Buczylowska & Petermann, 2018), it is not unlikely that age may account for more variability when including samples spanning multiple developmental periods. However, evidence so far suggests that during early-to-mid-childhood, age may not be the best marker of neural development of cognitive control. We show here that performance seems an adequate marker of neural maturity and processing of cognitive demand in several MD ss-fROIs during childhood.

Lastly, we explored whether MD ss-fROIs are already functionally distinct from the adjacent language network. In adults, the MD and language networks are differentiated in both function (Fedorenko et al., 2013) and resting-state connectivity (Blank, Kanwisher, & Fedorenko, 2014), but to date no one had explored whether this distinction is evident in child samples. We see greater selectivity for cognitive load compared to language in regions that were functionally localized using the SWM task (i.e., MD ss-fROIs) and we see greater within-than between-network resting-state connectivity for these same MD ss-fROIs compared to language ss-fROIs (functionally localized using the language localizer task). Both approaches suggest that even in kids as young as four years of age, we see functional distinction between the brain regions recruited for processing cognitive load versus language. We also see that within-MD network connectivity increases with age in the right hemisphere, but difference in connectivity does not change with age. Suggesting that these two networks are already dissociated at an early age, even though within network connectivity continues to increase.

Surprisingly, children show consistently higher activation in MD ss-fROIs in response to scrambled sentences than in response to typical sentences. This pattern is also observed in many of the adults’ MD ss-fROIs, which replicates a general pattern observed in an independent sample of adults (Diachek, Blank, Siegelman, Affourtit, & Fedorenko, 2020). This suggests that MD regions can be activated in a variety of situations when individuals engage with cognitively demanding stimuli (e.g., trying to unscramble sentences).

A few limitations should be addressed regarding the present study. First, we did not look at the association with out-of-scanner working memory performance, and therefore our findings may be limited in generalizability due to influences of the scanning environment. Doebel (2020) proposes that instead of being an integration of three separable but overlapping components (i.e., working memory, respond inhibition, shifting), EF is developed by integrating experience-based mental content (e.g., beliefs, knowledge, values) in service towards fulfilling specific goals. From this perspective, the capacity for children to self-regulate across contexts in service of goal-directed tasks increases throughout development (Perone, Simmering, & Buss, 2021). This suggests that neural selectivity observed in this study may not reflect the same neural network recruited by children in more naturalistic situations, especially for children who do not yet exhibit fully mature self-regulation capacity. Second, substantial motion is observed in child samples, which can produce spurious correlations in fMRI (Power et al., 2012). However, we accounted for this by implementing strict motion cutoffs in our cross-sectional sample, adding framewise displacement as a covariate for our longitudinal analyses, and comparing children to a motion matched adult group, who still show robust activation in MD ss-fROIs. Therefore, while the differences observed are likely due to maturational differences, and not a function of motion, they may not generalize to all children (i.e., those who exhibit higher motion during scans). Third, our child sample engaged in age-appropriate versions of the SWM task (to equate for task difficulty across age), thus it is possible that differences may be driven by difficulty level of the task. However, we see similar accuracy (during Hard trials) across age in our child sample, suggesting that the difficulty of the tasks was appropriate for engaging the MD ss-fROIs even for younger ages. Additionally, prior findings show that the MD network can be activated even by simple tasks in adults (Fedorenko et al., 2013; Wen, Mitchell, & Duncan, 2018), increasing confidence that differences in selectivity between children and adults were not driven by difficulty of the task. Future studies can explore how neural recruitment of MD ss-fROIs changes over time in a larger sample that allows evaluation of moderating and mediating variables. Further investigations that probe how neural markers relate to out of scanner behavior is warranted, so that we can be confident that these findings generalize across contexts.

In conclusion, this study provides evidence that a common neural network underlies processing cognitive demand in both children and adults. Though children do not yet exhibit fully mature selectivity of the MD network, they show adult-like patterns and performance related increases in MD selectivity in a sample of children as young as four years. These data further our understanding about the neural development underlying development of EF and how it relates to behavior. Identifying functional architecture that supports EF across development and typical variability furthers our understanding of executive dysfunction and early markers of risk. These findings can inform intervention research for individuals struggling with cognitive deficits.

## Acknowledgements

We would like to thank members of the Saygin Developmental Cognitive Neuroscience Lab for data collection. We would like to acknowledge the support from Center for Cognitive and Behavioral Brain Imaging (CCBBI) and Ohio Supercomputer Center (OSC). Z.M.S. was supported by the Alfred P. Sloan Foundation, OSU’s College of Arts & Sciences, and the Chronic Brain Injury initiative at OSU.

## Notes

**Conflict of Interest:** The authors declare no competing financial interests.

### Competing Interest Statement

The authors have declared no competing interest.

