## Supplemental Figure 1 for "Individual variability in performance reflects selectivity of the multiple demand network among children and adults"

*Depiction of spatial working memory task*

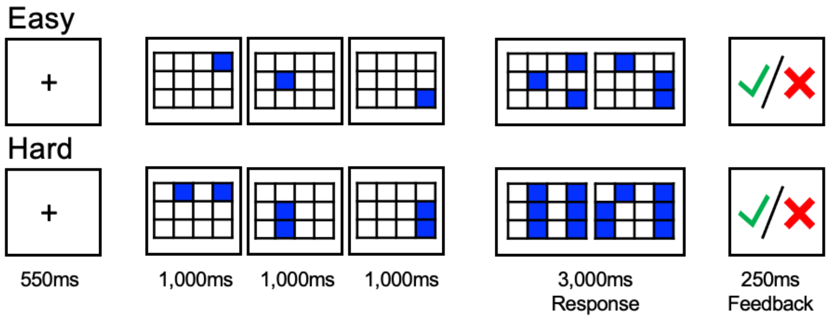


**Supplementary Figure 1.** Procedure and timing for the spatial working memory task (high difficulty) used to localize multiple demand network functional regions of interest.
