## Supplemental Table 4 for "Individual variability in performance reflects selectivity of the multiple demand network among children and adults"

***MD selectivity is associated with performance***

*Longitudinal sample*

We see a significant positive association between accuracy and MD selectivity in the "low" motion longitudinal child sample (**Supplemental Table 3**).

| | **Hard trials** | | **Easy trials** | |
|---|---|---|---|---|
| | **B** | **Main effect t-value** | **B** | **Main effect t-value** |
| **Right inferior frontal gyrus** | | | | |
| Accuracy | 2.71 | 2.28 | 3.66 | 9.44*** |
| Age | -0.41 | -3.71* | -0.45 | -13.32*** |
| Framewise displacement | 0.01 | 2.83* | $6.30 \times 10^{-3}$ | 10.02*** |
| **Right orbital middle frontal gyrus** | | | | |
| Accuracy | 4.02 | 2.03 | 5.22 | 4.24** |
| Age | -0.37 | 1.99 | -0.42 | -3.73** |
| Framewise displacement | 0.01 | 2.77* | 0.01 | 4.96** |
| **Right middle frontal gyrus** | | | | |
| Accuracy | 3.94 | 3.36* | 2.66 | 2.03 |
| Age | -0.46 | -4.03* | -0.32 | -2.86* |
| Framewise displacement | 0.01 | 2.68* | $3.15 \times 10^{-3}$ | 1.48 |
| **Right superior frontal sulcus** | | | | |
| Accuracy | *ns* | *ns* | 2.75 | 3.13* |
| Age | *ns* | *ns* | -0.20 | -2.50* |
| Framewise displacement | *ns* | *ns* | $3.50 \times 10^{-3}$ | 2.45* |
| **Right inferior frontal sulcus** | | | | |
| Accuracy | *ns* | *ns* | 2.80 | 3.97** |
| Age | *ns* | *ns* | -0.25 | -4.08** |
| Framewise displacement | *ns* | *ns* | $4.55 \times 10^{-3}$ | 3.97** |
| **Left posterior parietal** | | | | |
| Accuracy | 3.14 | 3.57* | *ns* | *ns* |
| Age | -0.41 | -4.74** | *ns* | *ns* |
| Framewise displacement | 0.01 | 2.95* | *ns* | *ns* |
| **Left superior parietal** | | | | |
| | *ns* | *ns* | 4.50 | 2.95* |
| | *ns* | *ns* | -0.45 | -2.91* |
| | *ns* | *ns* | $5.21 \times 10^{-3}$ | 2.05 |
| **Left precentral gyrus** | | | | |
| Accuracy | *ns* | *ns* | 3.03 | 1.95 |

| | | | | |
|---|---|---|---|---|
| Age | *ns* | *ns* | -0.43 | -2.71* |
| Framewise displacement | *ns* | *ns* | $4.29 \times 10^{-3}$ | 1.66 |
| **Left inferior frontal gyrus** | | | | |
| Accuracy | *ns* | *ns* | 2.93 | 2.23 |
| Age | *ns* | *ns* | -0.33 | -2.51* |
| Framewise displacement | *ns* | *ns* | $4.13 \times 10^{-3}$ | 1.89 |
| **Left anterior cingulate** | | | | |
| Accuracy | *ns* | *ns* | 3.40 | 4.21** |
| Age | *ns* | *ns* | -0.21 | -3.07* |
| Framewise displacement | *ns* | *ns* | $4.65 \times 10^{-3}$ | 3.52** |

**Supplemental Table 3.** Subject-specific fROIs that show significant main effects of accuracy, age, and/or framewise displacement in linear mixed effects models; “low motion” longitudinal child sample who also have complete data for all variables of interest at both scans (n = 6). Asterisks indicate strength of significance (uncorrected): * $p < .05$; ** $p < .01$; *** $p < .001$; *ns* = p > .05.

In the full sample of longitudinal children (n = 14; no motion cutoffs and some missing behavioral data), for models that account for variance explained by age and motion, we see fewer significant associations between accuracy during hard conditions and MD selectivity (**Supplemental Table 4**); in these models, motion was the most robust predictor of MD selectivity.

| | Hard trials | | Easy trials | |
|---|---|---|---|---|
| | B | Main effect t-value | B | Main effect t-value |
| **Right superior frontal sulcus** | | | | |
| Accuracy | 0.35 | 0.69 | 0.58 | 1.08 |
| Age | 0.02 | 0.28 | $7.33 \times 10^{-3}$ | 0.11 |
| Framewise displacement | $2.01 \times 10^{-3}$ | 3.56** | $2.08 \times 10^{-3}$ | 3.70** |
| **Right orbital middle frontal gyrus** | | | | |
| Accuracy | 0.73 | 1.03 | 0.22 | 0.29 |
| Age | -0.10 | -1.05 | -0.06 | -0.63 |
| Framewise displacement | $1.92 \times 10^{-3}$ | 2.45* | $1.84 \times 10^{-3}$ | 2.26* |
| **Right middle frontal gyrus** | | | | |
| Accuracy | 0.92 | 1.53 | 0.82 | 1.26 |
| Age | 0.02 | 0.27 | 0.04 | 0.54 |

| | | | | |
|---|---|---|---|---|
| Framewise displacement | $1.67x10^{-3}$ | 2.47* | $1.70x10^{-3}$ | 2.46* |
| **Left inferior parietal** | | | | |
| Accuracy | -1.02 | -1.43 | *ns* | *ns* |
| Age | 0.22 | 2.32* | *ns* | *ns* |
| Framewise displacement | $7.13x10^{-3}$ | 0.90 | *ns* | *ns* |
| **Right posterior parietal** | | | | |
| Accuracy | 0.16 | 0.29 | 0.28 | 0.47 |
| Age | -0.07 | -0.89 | -0.08 | - 1.01 |
| Framewise displacement | $1.50x10^{-3}$ | 2.33* | $1.58x10^{-3}$ | 2.45* |

**Supplemental Table 4.** Subject-specific fROIs that show significant main effects of age or framewise displacement in linear mixed effects models for the full longitudinal child sample (n = 14); no motion cutoffs were applied, and some subjects are missing accuracy data at one timepoint. Asterisks indicate strength of significance (uncorrected): $^{t}$ p < .07; * *p* < .05; ** p < .01; *** p < .001; ns = p > .05.
