## Supplemental Table 2 for "Individual variability in performance reflects selectivity of the multiple demand network among children and adults"

***MD selectivity is associated with performance***

*Cross-sectional*

Pearson's correlations between accuracy and selectivity of several MD ss-fROIs show a significant relationship during both Hard and Easy trials; significant correlations were primarily in the right hemisphere (**Supplemental Table 1**). Because there is a significant correlation between accuracy during easy trials and age, we also conducted multiple linear regression to assess the unique effect of accuracy on MD selectivity, while controlling for variability associated with age (**Table3**; **Supplemental Table 2**).

| | Hard trials | | Easy trials | |
|---|---|---|---|---|
| | t-value | Pearson's r | t-value | Pearson's r |
| **Right Hemisphere** | | | | |
| Posterior parietal | 1.56 | 0.27 | -0.38 | -0.07 |
| Superior parietal | 2.12* | 0.36 | 2.03 | 0.35 |
| Inferior parietal | 3.56** | $0.55^{m}$ | 1.82 | 0.32 |
| Precentral gyrus | 3.22** | $0.51^{m}$ | 2.18* | 0.37 |
| Superior frontal sulcus | 2.75** | $0.45^{m}$ | 2.81** | $0.46^{m}$ |
| Inferior frontal sulcus | 4.21*** | $0.61^{m}$ | 2.92** | $0.47^{m}$ |
| Middle frontal gyrus | 4.41*** | $0.63^{m}$ | 3.84*** | $0.57^{m}$ |
| Orbital middle frontal gyrus | 1.83 | 0.32 | 1.66 | 0.29 |
| Inferior frontal gyrus | 1.18 | 0.21 | 1.12 | 0.20 |
| Anterior cingulate cortex | 2.20* | 0.37 | 3.44** | $0.53^{m}$ |
| **Left Hemisphere** | | | | |
| Posterior parietal | 2.66* | $0.44^{m}$ | 0.62 | 0.11 |
| Superior parietal | 2.42* | $0.40^{m}$ | 1.48 | 0.26 |
| Inferior parietal | 1.88 | 0.32 | 1.42 | 0.25 |
| Precentral gyrus | 1.03 | 0.18 | -0.002 | -0.0003 |
| Superior frontal sulcus | 2.18* | 0.37 | 2.13* | 0.36 |
| Inferior frontal sulcus | 1.60 | 0.28 | 2.21* | 0.37 |
| Middle frontal gyrus | 1.77 | 0.31 | 1.87 | 0.32 |
| Orbital middle frontal gyrus | 0.67 | 0.12 | 1.17 | 0.21 |
| Inferior frontal gyrus | 1.20 | 0.21 | 2.46* | 0.41 |
| Anterior cingulate cortex | 1.99 | 0.34 | 2.26* | 0.38 |

**Supplemental Table 1.** Pearson correlations between accuracy during Hard and Easy trials and selectivity of MD ss-fROIs. Asterisks indicate strength of significance (uncorrected): * $p < .05$; ** $p < .01$; *** $p < .001$; $^{m}$ Bonferroni-Holm corrected $p < .05$ (3 parietal regions, 7 frontal regions).

| | Accuracy B | Accuracy t-value | Age B | Age t-value | Full model F-value | Adjusted $R^2$ |
|---|---|---|---|---|---|---|
| **Right Hemisphere** | | | | | | |
| Posterior parietal | 0.17 | 0.46 | -0.06 | -1.84 | 1.78 | 0.05 |
| Superior parietal | 1.02 | 2.21* | -0.04 | -0.89 | 2.44 | 0.09 |
| Inferior parietal | 0.85 | 2.17 | -0.04 | -1.19 | 2.38 | 0.08 |
| Precentral gyrus | 0.82 | 2.05* | -0.01 | -0.28 | 2.35 | 0.08 |
| Superior frontal sulcus | 1.21 | 2.78** | -0.03 | -0.64 | 4.08* | 0.17 |
| Inferior frontal sulcus | 0.85 | 2.45* | 0.01 | 0.32 | 4.20* | 0.17 |
| Middle frontal gyrus | 1.45 | 3.56** | -0.01 | -0.35 | 7.22** | 0.29 [m] |
| Orbital middle frontal gyrus | 0.54 | 1.05 | 0.05 | 1.03 | 1.92 | 0.06 |
| Inferior frontal gyrus | 0.35 | 0.82 | 0.02 | 0.40 | 0.68 | -0.02 |
| Anterior cingulate cortex | 1.13 | 2.57* | 0.05 | 1.27 | 6.86** | 0.27 [m] |
| **Left Hemisphere** | | | | | | |
| Posterior parietal | 0.47 | 1.11 | -0.05 | -1.25 | 0.98 | -0.002 |
| Superior parietal | 0.79 | 1.53 | -0.02 | -0.50 | 1.19 | 0.01 |
| Inferior parietal | 0.56 | 1.12 | 0.01 | 0.32 | 1.03 | 0.002 |
| Precentral gyrus | 0.05 | 0.13 | -0.01 | -0.30 | 0.05 | -0.07 |
| Superior frontal sulcus | 0.94 | 1.93 | $4.72 \times 10^{-3}$ | 0.11 | 2.19 | 0.07 |
| Inferior frontal sulcus | 0.77 | 1.46 | 0.06 | 1.24 | 3.25 | 0.13 |
| Middle frontal gyrus | 0.76 | 1.49 | 0.02 | 0.39 | 1.78 | 0.05 |
| Orbital middle frontal gyrus | 0.30 | 0.59 | 0.05 | 1.04 | 1.23 | 0.01 |
| Inferior frontal gyrus | 0.68 | 1.61 | 0.06 | 1.45 | 4.18* | 0.17 |
| Anterior cingulate cortex | 0.71 | 1.49 | 0.06 | 1.27 | 3.41 | 0.14 |

**Supplemental Table 2.** Main effect of accuracy during Easy trials on selectivity of MD ss-fROIs during a spatial working memory task for our child sample. Multiple linear regressions were conducted to account for variability associated with age. Asterisks indicate strength of significance (uncorrected): * $p < .05$; ** $p < .01$; *** $p < .001$; [m] Bonferroni-Holm full model corrected $p < .05$ (3 parietal regions, 7 frontal regions).
