## Supplemental Figure 2 for "Individual variability in performance reflects selectivity of the multiple demand network among children and adults"

***Behavioral metrics during SWM task positively associated with age***

In the cross-sectional sample, we see only a significant correlation between age and performance during easy trials. Children show higher accuracy and quicker reaction times with increasing age. However, these relationships do not emerge during hard trials. This may suggest that though age is a good predictor of how well children are able to engage in easy trials of this task, other variables are likely a better predictor of how well children perform in the face of more cognitively demanding tasks.

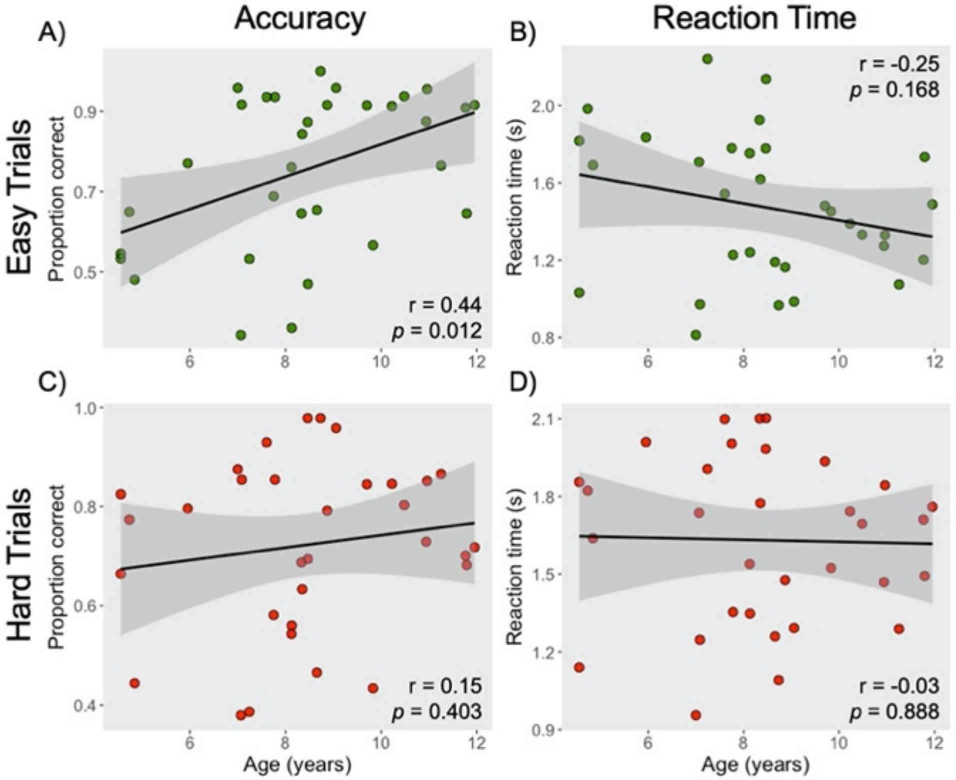


**Supplemental Figure 2.** Pearson correlations for age and behavioral metrics during the spatial working memory task in the cross-sectional child sample. Accuracy (A,C) and reaction time (B,D) during Easy (green) and Hard (red) trials are reported separately.
